## Supplementary material for "PyLandStats: An open-source Pythonic library to compute landscape metrics": Text S1 - PyLandStats Documentation

***Release 1.0.2***

**Martí Bosch**

**Jul 25, 2019**

### REFERENCE GUIDE:


1 Landscape analysis 3

2 Spatiotemporal analysis 11

3 Gradient analysis 13

4 Spatiotemporal buffer analysis 17

5 Change log 21

6 Contributing 25

7 Features 27

8 Using PyLandStats 29

9 Indices and tables 31

Index 33

Open-source Pythonic library to compute landscape metrics within the PyData stack (NumPy, pandas, matplotlib. . . )

CHAPTER

ONE

### LANDSCAPE ANALYSIS

#### 1.1 List of implemented metrics

The metrics of PyLandStats are computed according to its definitions in FRAGSTATS.

The notation for the metrics below is as follows:

- the letters with suffixes $a_{i,j}, p_{i,j}, h_{i,j}$ respectively represent the area, perimeter, and distance to the nearest neighboring patch of the same class of the patch $j$ of class $i$.
- the letters with suffixes $e_{i,k}, g_{i,k}$ respectively represent the total edge between and number of pixel adjacencies between classes $i$ and $k$
- the capital letters $A, N, E$ respectively represent the total area, total number of patches and total edge of the landscape

Like FRAGSTATS, PyLandStats features six distribution-statistics metrics for each patch-level metric, which consist in a statistical aggregation of the values computed for each patch of a class or the whole landscape:

- the mean, which can be computed by adding a *_mn* suffix to the method name, e.g., *area_mn*
- the area-weighted mean, which can be computed by adding a *_am* suffix to the method name, e.g., *area_am*
- the median, which can be computed by adding a *_md* suffix to the method name, e.g., *area_md*
- the range, which can be computed by adding a *_ra* suffix to the method name, e.g., *area_ra*
- the standard deviation, which can be computed by adding a *_sd* suffix to the method name, e.g., *area_sd*
- the coefficient of variation, which can be computed by adding a *_cv* suffix to the method name, e.g., *area_cv*

note that the distribution-statistics metrics do not appear in the documentation below.

See the FRAGSTATS documentation for more information.

##### 1.1.1 Patch-level metrics

###### Area, density, edge

`Landscape.`**`area`**`(`*class_val=None*, *hectares=True*`)`

The area of each patch of the landscape

$$AREA = a_{i,j} \quad [hec] \; or \; [m]$$

**Parameters**

- **class_val** (*int, optional*) – If provided, the metric will be computed for the corresponding class only, otherwise it will be computed for all the classes of the landscape
- **hectares** (*bool, default True*) – Whether the landscape area should be converted to hectares (tends to yield more legible values for the metric)

**Returns** **AREA** – AREA > 0, without limit

**Return type** pd.Series if *class_val* is provided, pd.DataFrame otherwise

`Landscape.`**`perimeter`**(*class_val=None*)

The perimeter of each patch of the landscape

$$PERIM = p_{i,j} \quad [m]$$

**Parameters** **class_val** (*int, optional*) – If provided, the metric will be computed for the corresponding class only, otherwise it will be computed for all the classes of the landscape

**Returns** **PERIM** – PERIM > 0, without limit

**Return type** pd.Series if *class_val* is provided, pd.DataFrame otherwise

#### Shape

`Landscape.`**`perimeter_area_ratio`**(*class_val=None*, *hectares=True*)

The ratio between the perimeter and area of each patch of the landscape. Measures shape complexity, however it varies with the size of the patch, e.g, for the same shape, larger patches will have a smaller perimeter-area ratio.

$$PARA = \frac{p_{i,j}}{a_{i,j}} \quad [m/hec] \; or \; [m/m^2]$$

**Parameters**

- **class_val** (*int, optional*) – If provided, the metric will be computed for the corresponding class only, otherwise it will be computed for all the classes of the landscape
- **hectares** (*bool, default True*) – Whether the area should be converted to hectares (tends to yield more legible values for the metric)

**Returns** **PARA** – PARA > 0, without limit

**Return type** pd.Series if *class_val* is provided, pd.DataFrame otherwise

`Landscape.`**`shape_index`**(*class_val=None*)

A measure of shape complexity, similar to the perimeter-area ratio, but correcting for its size problem by adjusting for a standard square shape.

$$SHAPE = \frac{.25 \; p_{i,j}}{\sqrt{a_{i,j}}}$$

**Parameters** **class_val** (*int, optional*) – If provided, the metric will be computed for the corresponding class only, otherwise it will be computed for all the classes of the landscape

**Returns** **SHAPE** – SHAPE >= 1, without limit ; SHAPE equals 1 when the patch is maximally compact, and increases without limit as patch shape becomes more regular

**Return type** pd.Series if *class_val* is provided, pd.DataFrame otherwise

`Landscape.`**`fractal_dimension`**(*class_val=None*)

A measure of shape complexity appropriate across a wide range of patch sizes

$$FRAC = \frac{2 \; ln(.25 \; p_{i,j})}{ln(a_{i,j})}$$

**Parameters** **class_val** (*int, optional*) – If provided, the metric will be computed for the corresponding class only, otherwise it will be computed for all the classes of the landscape

**Returns** **FRAC** – 1 <= FRAC <=2 ; for a two-dimensional patch, FRAC approaches 1 for very simple shapes such as squares, and approaches 2 for complex plane-filling shapes

**Return type** pd.Series if *class_val* is provided, pd.DataFrame otherwise

##### Aggregation

`Landscape.`**`euclidean_nearest_neighbor`**(*class_val=None*)

Distance to the nearest neighboring patch of the same class based on the shortest edge-to-edge distance

$$ENN = h_{i,j} \quad [m]$$

**Parameters** **class_val** (*int, optional*) – If provided, the metric will be computed for the corresponding class only, otherwise it will be computed for all the classes of the landscape

**Returns** **ENN** – ENN > 0, without limit ; ENN approaches 0 as the distance to the nearest neighbors decreases

**Return type** numeric

#### 1.1.2 Class-level and landscape-level metrics

##### Area, density, edge

`Landscape.`**`total_area`**(*class_val=None*, *hectares=True*)

Total area. If *class_val* is provided, the metric is computed at the class level as in:

$$TA_i = \sum_{j=1}^{n_i} a_{i,j} \quad [hec] \; or \; [m] \quad (class\ i)$$

otherwise, the metric is computed at the landscape level as in:

**..math::** TA = A quad [hec] ; or ; [m] quad (landscape)

**Parameters**

- **class_val** (*int, optional*) – If provided, the metric will be computed at the level of the corresponding class, otherwise it will be computed at the landscape level
- **hectares** (*bool, default True*) – Whether the area should be converted to hectares (tends to yield more legible values for the metric)

**Returns** **TA**

**Return type** numeric

`Landscape.`**`proportion_of_landscape`**(*class_val*, *percent=True*)

Measures the proportional abundance of a particular class within the landscape. It is computed at the class level as in:

$$PLAND = \frac{1}{A} \sum_{j}^{n_i} a_{i,j}$$

**Parameters**

- **class_val** (*int*) – Class for which the metric should be computed
- **percent** (*bool, default True*) – Whether the index should be expressed as proportion or converted to percentage. If True, this method returns FRAGSTATS' percentage of landscape (PLAND)

**Returns PLAND** – 0 < PLAND <= 100 ; PLAND approaches 0 when the occurrence of the corresponding class becomes increasingly rare, and approaches 100 when the entire landscape consists of a single patch of such class.

**Return type** numeric

Landscape.**number_of_patches**(*class_val=None*)

Number of patches. If *class_val* is provided, the metric is computed at the class level as in:

$$NP_i = n_i \quad (class\ i)$$

otherwise, the metric is computed at the landscape level as in:

$$NP = N \quad (landscape)$$

**Parameters class_val** (*int, optional*) – If provided, the metric will be computed at the level of the corresponding class, otherwise it will be computed at the landscape level

**Returns NP** – NP >= 1, without limit

**Return type** int

Landscape.**patch_density**(*class_val=None*, *percent=True*, *hectares=True*)

Density of class patches, which is arguably more useful than the number of patches since it facilitates comparison among landscapes of different sizes. If *class_val* is provided, the metric is computed at the class level as in:

$$PD_i = \frac{n_i}{A} \quad [1/hec]\ or\ [1/m^2] \quad (class\ i)$$

otherwise, the metric is computed at the landscape level as in:

$$PD = \frac{N}{A} \quad [1/hec]\ or\ [1/m^2] \quad (landscape)$$

**Parameters**

- **class_val** (*int, optional*) – If provided, the metric will be computed at the level of the corresponding class, otherwise it will be computed at the landscape level
- **percent** (*bool, default True*) – Whether the index should be expressed as proportion or converted to percentage
- **hectares** (*bool, default True*) – Whether the landscape area should be converted to hectares (tends to yield more legible values for the metric)

**Returns PD** – PD > 0, constrained by cell size ; maximum PD is attained when every cell is a separate patch

**Return type** numeric

Landscape.**largest_patch_index**(*class_val=None*, *percent=True*)

The proportion of total landscape comprised by the largest patch. If *class_val* is provided, the metric is computed at the class level as in:

$$LPI_i = \frac{1}{A} \max_{j=1}^{n_i} a_{i,j} \quad (class\ i)$$

otherwise, the metric is computed at the landscape level as in:

$$LPI = \frac{1}{A} \max a_{i,j} \quad (landscape)$$

**Parameters**

- **class_val** (*int, optional*) – If provided, the metric will be computed at the level of the corresponding class, otherwise it will be computed at the landscape level
- **percent** (*bool, default True*) – Whether the index should be expressed as proportion or converted to percentage

**Returns** **LPI** – 0 < LPI <= 100 (or 0 < LPI <= 1 if percent argument is False) ; LPI approaches 0 when the largest patch of the corresponding class is increasingly small, and approaches its maximum value when such largest patch comprises the totality of the landscape

**Return type** numeric

`Landscape.total_edge`(*class_val=None*, *count_boundary=False*)

Measure of the total edge length. If *class_val* is provided, the metric is computed at the class level as in:

$$TE_i = \sum_{k=1}^{m} e_{i,k} \quad [m] \quad (class\ i)$$

otherwise, the metric is computed at the landscape level as in:

$$TE = E \quad [m] \quad (landscape)$$

**Parameters**

- **class_val** (*int, optional*) – If provided, the metric will be computed at the level of the corresponding class, otherwise it will be computed at the landscape level
- **count_boundary** (*bool, default False*) – Whether the boundary of the landscape should be included in the total edge length

**Returns** **TE** – TE >= 0 ; TE equals 0 when the entire landscape and its border consist of the corresponding class

**Return type** numeric

`Landscape.edge_density`(*class_val=None*, *count_boundary=False*, *hectares=True*)

Measure of edge length per area unit, which facilitates comparison among landscapes of different sizes. If *class_val* is provided, the metric is computed at the class level as in:

$$ED_i = \frac{1}{A} \sum_{k=1}^{m} e_{i,k} \quad [m/hec]\ or\ [m/m^2] \quad (class\ i)$$

otherwise, the metric is computed at the landscape level as in:

$$ED = \frac{E}{A} \quad [m/hec]\ or\ [m/m^2] \quad (landscape)$$

**Parameters**

- **class_val** (*int, optional*) – If provided, the metric will be computed at the level of the corresponding class, otherwise it will be computed at the landscape level
- **count_boundary** (*bool, default False*) – Whether the boundary of the landscape should be considered
- **hectares** (*bool, default True*) – Whether the landscape area should be converted to hectares (tends to yield more legible values for the metric)

**Returns** **ED** – ED >= 0, without limit ; ED equals 0 when the entire landscape and its border consist of the corresponding patch class.

**Return type** numeric

##### Aggregation

`Landscape.`**`landscape_shape_index`**(*class_val=None*)

Measure of class aggregation that provides a standardized measure of edginess that adjusts for the size of the landscape. If *class_val* is provided, the metric is computed at the class level as in:

$$LSI_i = \frac{.25 \sum\limits_{k=1}^{m} e_{i,k}}{\sqrt{A}} \quad (class\ i)$$

otherwise, the metric is computed at the landscape level as in:

$$LSI = \frac{.25E}{\sqrt{A}} \quad (landscape)$$

**Parameters** **`class_val`** (*`int, optional`*) – If provided, the metric will be computed at the level of the corresponding class, otherwise it will be computed at the landscape level

**Returns** **LSI** – LSI >=1 ; LSI equals 1 when the entire landscape consists of a single patch of the corresponding class, and increases without limit as the patches of such class become more disaggregated.

**Return type** float

#### 1.1.3 Landscape-level metrics

##### Contagion, interspersion

`Landscape.`**`contagion`**(*percent=True*)

Measure of aggregation that measures the probability that two random adjacent cells belong to the same class. It is computed at the landscape level as in:

$$CONTAG = 1 + \frac{\sum\limits_{i=1}^{m} \sum\limits_{k=1}^{m} \left[ P_i \frac{g_{i,k}}{\sum\limits_{k=1}^{m} g_{i,k}} \right] \left[ ln \left( P_i \frac{g_{i,k}}{\sum\limits_{k=1}^{m} g_{i,k}} \right) \right]}{2ln(m)}$$

**Parameters** **`percent`** (*`bool, default True`*) – Whether the index should be expressed as proportion or converted to percentage

**Returns** **CONTAG** – 0 < CONTAG <= 100 ; CONTAG approaches 0 when the classes are maximally disaggregated (i.e., every cell is a patch of a different class) and interspersed (i.e., equal proportions of all pairwise adjacencies), and approaches its maximum when the landscape consists of a single patch.

**Return type** float

`Landscape.`**`shannon_diversity_index`**()

Measure of diversity that reflects the number of classes present in the landscape as well as the relative abundance of each class. It is computed at the landscape level as in:

$$SHDI = -\sum_{i=1}^{m} \left( P_i \ lnP_i \right)$$

**Returns** **SHDI** – SHDI >= 0 ; SHDI approaches 0 when the entire landscape consists of a single patch, and increases as the number of classes increases and/or the proportional distribution of area among classes becomes more equitable.

**Return type** float

#### 1.2 Computing metrics data frames

`Landscape.`**`compute_patch_metrics_df`**(*metrics=None*, *metrics_kws={}*)

Computes the patch-level metrics

**Parameters**

- **metrics** (*list-like, optional*) – A list-like of strings with the names of the metrics that should be computed. If None, all the implemented patch-level metrics will be computed.
- **metrics_kws** (*dict, optional*) – Dictionary mapping the keyword arguments (values) that should be passed to each metric method (key), e.g., to compute *area* in meters instead of hectares, metric_kws should map the string 'area' (method name) to {'hectares': False}. The default empty dictionary will compute each metric according to FRAGSTATS defaults.

**Returns** **df** – Dataframe with the values computed for each patch (index) and metric (columns)

**Return type** pd.DataFrame

`Landscape.`**`compute_class_metrics_df`**(*metrics=None*, *metrics_kws={}*)

Computes the class-level metrics

**Parameters**

- **metrics** (*list-like, optional*) – A list-like of strings with the names of the metrics that should be computed. If None, all the implemented class-level metrics will be computed.
- **metrics_kws** (*dict, optional*) – Dictionary mapping the keyword arguments (values) that should be passed to each metric method (key), e.g., to exclude the boundary from the computation of *total_edge*, metric_kws should map the string 'total_edge' (method name) to {'count_boundary': False}. The default empty dictionary will compute each metric according to FRAGSTATS defaults.

**Returns** **df** – Dataframe with the values computed for each class (index) and metric (columns)

**Return type** pd.DataFrame

`Landscape.`**`compute_landscape_metrics_df`**(*metrics=None*, *metrics_kws={}*)

Computes the landscape-level metrics

**Parameters**

- **metrics** (*list-like, optional*) – A list-like of strings with the names of the metrics that should be computed. If None, all the implemented landscape-level metrics will be computed.
- **metrics_kws** (*dict, optional*) – Dictionary mapping the keyword arguments (values) that should be passed to each metric method (key), e.g., to exclude the boundary from the computation of *total_edge*, metric_kws should map the string 'total_edge' (method name) to {'count_boundary': False}. The default empty dictionary will compute each metric according to FRAGSTATS defaults.

**Returns** **df** – Dataframe with the values computed at the landscape level (one row only) for each metric (columns)

**Return type** pd.DataFrame

#### 1.3 Plotting landscape raster

`Landscape.`**`plot_landscape`**(*cmap=None*, *ax=None*, *legend=False*, *figsize=None*, ***show_kws*)

Plots the landscape with a categorical legend by means of *rasterio.plot.show*

**Parameters**

- **`cmap`** (str or *~matplotlib.colors.Colormap*, optional) – A Colormap instance
- **`ax`** (*`axis object, optional`*) – Plot in given axis; if None creates a new figure
- **`legend`** (*`bool, optional`*) – If `True`, display the legend
- **`figsize`** (*`tuple of two numeric types, optional`*) – Size of the figure to create. Ignored if axis *ax* is provided
- **`**show_kws`** (*`optional`*) – Keyword arguments to be passed to *rasterio.plot.show*

**Returns** **ax** – axis with plot data

**Return type** matplotlib axis

CHAPTER

TWO

### SPATIOTEMPORAL ANALYSIS

**class** pylandstats.**SpatioTemporalAnalysis**(*landscapes*, *metrics=None*, *classes=None*, *dates=None*, *metrics_kws={}*)

**__init__**(*landscapes*, *metrics=None*, *classes=None*, *dates=None*, *metrics_kws={}*)

**Parameters**

- **landscapes** (*list-like*) – A list-like of *Landscape* objects or of strings/file objects/ pathlib.Path objects so that each is passed as the *landscape* argument of *Landscape.__init__*
- **metrics** (*list-like, optional*) – A list-like of strings with the names of the metrics that should be computed in the context of this analysis case
- **classes** (*list-like, optional*) – A list-like of ints or strings with the class values that should be considered in the context of this analysis case
- **dates** (*list-like, optional*) – A list-like of ints or strings that label the date of each snapshot of *landscapes* (for DataFrame indices and plot labels)
- **metrics_kws** (*dict, optional*) – Dictionary mapping the keyword arguments (values) that should be passed to each metric method (key), e.g., to exclude the boundary from the computation of *total_edge*, metric_kws should map the string 'total_edge' (method name) to {'count_boundary': False}. The default empty dictionary will compute each metric according to FRAGSTATS defaults.

**property class_metrics_df**

Property that computes the data frame of class-level metrics, which is multi-indexed by the class and date. Once computed, the data frame is cached so further calls to the property just access an attribute and therefore run in constant time.

**property landscape_metrics_df**

Property that computes the data frame of landcape-level metrics, which is indexed by the date. Once computed, the data frame is cached so further calls to the property just access an attribute and therefore run in constant time.

**plot_landscapes**(*cmap=None*, *legend=True*, *subplots_kws={}*, *show_kws={}*, *subplots_adjust_kws={}*)

Plots each landscape snapshot in a dedicated matplotlib axis by means of the *Landscape.plot_landscape* method of each instance

**Parameters**

- **cmap** (str or ~*matplotlib.colors.Colormap*, optional) – A Colormap instance
- **legend** (*bool, optional*) – If `True`, display the legend of the land use/cover color codes

- **subplots_kws** (*dict, optional*) – Keyword arguments to be passed to *plt.subplots*
- **show_kws** (*dict, optional*) – Keyword arguments to be passed to *rasterio.plot.show*
- **subplots_adjust_kws** (*dict, optional*) – Keyword arguments to be passed to *plt.subplots_adjust*

**Returns** **fig** – The figure with its corresponding plots drawn into its axes

**Return type** *matplotlib.figure.Figure*

**plot_metric**(*metric*, *class_val=None*, *ax=None*, *metric_legend=True*, *metric_label=None*, *fmt='–o'*, *plot_kws={}*, *subplots_kws={}*)

**Parameters**

- **metric** (*str*) – A string indicating the name of the metric to plot
- **class_val** (*int, optional*) – If provided, the metric will be plotted at the level of the corresponding class, otherwise it will be plotted at the landscape level
- **ax** (*axis object, optional*) – Plot in given axis; if None creates a new figure
- **metric_legend** (*bool, default True*) – Whether the metric label should be displayed within the plot (as label of the y-axis)
- **metric_label** (*str, optional*) – Label of the y-axis to be displayed if *metric_legend* is *True*. If the provided value is *None*, the label will be taken from the *settings* module
- **fmt** (*str, default '--o'*) – A format string for *plt.plot*
- **plot_kws** (*dict*) – Keyword arguments to be passed to *plt.plot*
- **subplots_kws** (*dict*) – Keyword arguments to be passed to *plt.subplots*, only if no axis is given (through the *ax* argument)

**Returns** **ax** – Returns the Axes object with the plot drawn onto it

**Return type** axis object

CHAPTER

THREE

### GRADIENT ANALYSIS

**class** pylandstats.**GradientAnalysis**(*landscape*, *masks_arr*, *attribute_name=None*, *attribute_values=None*, ***kwargs*)

**__init__**(*landscape*, *masks_arr*, *attribute_name=None*, *attribute_values=None*, ***kwargs*)

**Parameters**

- **landscapes** (*list-like*) – A list-like of *Landscape* objects or of strings/file objects/ pathlib.Path objects so that each is passed as the *landscape* argument of *Landscape.__init__*
- **masks_arr** (*list-like or np.ndarray*) – A list-like of numpy arrays of shape (width, height), i.e., of the same shape as the landscape raster image. Each array will serve to mask the base landscape and define a region of study for which the metrics will be computed separately. The same information can also be provided as a single array of shape (num_masks, width, height).
- **attribute_name** (*str, optional*) – Name of the attribute that will distinguish each landscape
- **attribute_values** (*str, optional*) – Values of the attribute that correspond to each of the landscapes

**property class_metrics_df**

Property that computes the data frame of class-level metrics, which is multi-indexed by the class and attribute value. Once computed, the data frame is cached so further calls to the property just access an attribute and therefore run in constant time.

**property landscape_metrics_df**

Property that computes the data frame of landscape-level metrics, which is indexed by the attribute value. Once computed, the data frame is cached so further calls to the property just access an attribute and therefore run in constant time.

**plot_landscapes**(*cmap=None*, *legend=True*, *subplots_kws={}*, *show_kws={}*, *subplots_adjust_kws={}*)

Plots each landscape snapshot in a dedicated matplotlib axis by means of the *Landscape.plot_landscape* method of each instance

**Parameters**

- **cmap** (str or *~matplotlib.colors.Colormap*, optional) – A Colormap instance
- **legend** (*bool, optional*) – If `True`, display the legend of the land use/cover color codes
- **subplots_kws** (*dict, optional*) – Keyword arguments to be passed to *plt.subplots*

- **show_kws** (*dict, optional*) – Keyword arguments to be passed to *rasterio.plot.show*
- **subplots_adjust_kws** (*dict, optional*) – Keyword arguments to be passed to *plt.subplots_adjust*

**Returns** **fig** – The figure with its corresponding plots drawn into its axes

**Return type** *matplotlib.figure.Figure*

**plot_metric**(*metric*, *class_val=None*, *ax=None*, *metric_legend=True*, *metric_label=None*, *fmt='–o'*, *plot_kws={}*, *subplots_kws={}*)

**Parameters**

- **metric** (*str*) – A string indicating the name of the metric to plot
- **class_val** (*int, optional*) – If provided, the metric will be plotted at the level of the corresponding class, otherwise it will be plotted at the landscape level
- **ax** (*axis object, optional*) – Plot in given axis; if None creates a new figure
- **metric_legend** (*bool, default True*) – Whether the metric label should be displayed within the plot (as label of the y-axis)
- **metric_label** (*str, optional*) – Label of the y-axis to be displayed if *metric_legend* is *True*. If the provided value is *None*, the label will be taken from the *settings* module
- **fmt** (*str, default '--o'*) – A format string for *plt.plot*
- **plot_kws** (*dict*) – Keyword arguments to be passed to *plt.plot*
- **subplots_kws** (*dict*) – Keyword arguments to be passed to *plt.subplots*, only if no axis is given (through the *ax* argument)

**Returns** **ax** – Returns the Axes object with the plot drawn onto it

**Return type** axis object

**class** pylandstats.**BufferAnalysis**(*landscape*, *base_mask*, *buffer_dists*, *buffer_rings=False*, *base_mask_crs=None*, *landscape_crs=None*, *landscape_transform=None*, *metrics=None*, *classes=None*, *metrics_kws={}*)

**__init__**(*landscape*, *base_mask*, *buffer_dists*, *buffer_rings=False*, *base_mask_crs=None*, *landscape_crs=None*, *landscape_transform=None*, *metrics=None*, *classes=None*, *metrics_kws={}*)

**Parameters**

- **landscapes** (*list-like*) – A list-like of *Landscape* objects or of strings/file objects/ pathlib.Path objects so that each is passed as the *landscape* argument of *Landscape.__init__*
- **base_mask** (*shapely geometry or geopandas GeoSeries*) – Geometry that will serve as a base mask to buffer around
- **buffer_dists** (*list-like*) – Buffer distances
- **buffer_rings** (*bool, default False*) – If *False*, each buffer zone will consist of the whole region that lies within the respective buffer distance around the base mask. If *True*, buffer zones will take the form of rings around the base mask.

- **base_mask_crs** (*dict, optional*) – The coordinate reference system of the base mask. Required if the base mask is a shapely geometry or a geopandas GeoSeries without the *crs* attribute set
- **landscape_crs** (*dict, optional*) – The coordinate reference system of the landscapes. Required if the passed-in landscapes are *Landscape* objects, ignored if they are paths to GeoTiff rasters that already contain such information.
- **landscape_transform** (*affine.Affine*) – Transformation from pixel coordinates to coordinate reference system. Required if the passed-in landscapes are *Landscape* objects, ignored if they are paths to GeoTiff rasters that already contain such information.
- **metrics** (*list-like, optional*) – A list-like of strings with the names of the metrics that should be computed in the context of this analysis case
- **classes** (*list-like, optional*) – A list-like of ints or strings with the class values that should be considered in the context of this analysis case
- **metrics_kws** (*dict, optional*) – Dictionary mapping the keyword arguments (values) that should be passed to each metric method (key), e.g., to exclude the boundary from the computation of *total_edge*, metric_kws should map the string 'total_edge' (method name) to {'count_boundary': False}. The default empty dictionary will compute each metric according to FRAGSTATS defaults.

**property class_metrics_df**

Property that computes the data frame of class-level metrics, which is multi-indexed by the class and buffer distance. Once computed, the data frame is cached so further calls to the property just access an attribute and therefore run in constant time.

**property landscape_metrics_df**

Property that computes the data frame of landcape-level metrics, which is indexed by the buffer distance. Once computed, the data frame is cached so further calls to the property just access an attribute and therefore run in constant time.

**plot_landscapes**(*cmap=None*, *legend=True*, *subplots_kws={}*, *show_kws={}*, *subplots_adjust_kws={}*)

Plots each landscape snapshot in a dedicated matplotlib axis by means of the *Landscape.plot_landscape* method of each instance

**Parameters**

- **cmap** (str or *~matplotlib.colors.Colormap*, optional) – A Colormap instance
- **legend** (*bool, optional*) – If `True`, display the legend of the land use/cover color codes
- **subplots_kws** (*dict, optional*) – Keyword arguments to be passed to *plt.subplots*
- **show_kws** (*dict, optional*) – Keyword arguments to be passed to *rasterio.plot.show*
- **subplots_adjust_kws** (*dict, optional*) – Keyword arguments to be passed to *plt.subplots_adjust*

**Returns** **fig** – The figure with its corresponding plots drawn into its axes

**Return type** *matplotlib.figure.Figure*

**plot_metric**(*metric*, *class_val=None*, *ax=None*, *metric_legend=True*, *metric_label=None*, *fmt='–o'*, *plot_kws={}*, *subplots_kws={}*)

**Parameters**

- **metric** (*str*) – A string indicating the name of the metric to plot
- **class_val** (*int, optional*) – If provided, the metric will be plotted at the level of the corresponding class, otherwise it will be plotted at the landscape level
- **ax** (*axis object, optional*) – Plot in given axis; if None creates a new figure
- **metric_legend** (*bool, default True*) – Whether the metric label should be displayed within the plot (as label of the y-axis)
- **metric_label** (*str, optional*) – Label of the y-axis to be displayed if *metric_legend* is *True*. If the provided value is *None*, the label will be taken from the *settings* module
- **fmt** (*str, default '--o'*) – A format string for *plt.plot*
- **plot_kws** (*dict*) – Keyword arguments to be passed to *plt.plot*
- **subplots_kws** (*dict*) – Keyword arguments to be passed to *plt.subplots*, only if no axis is given (through the *ax* argument)

**Returns** **ax** – Returns the Axes object with the plot drawn onto it

**Return type** axis object

CHAPTER

FOUR

### SPATIOTEMPORAL BUFFER ANALYSIS

**class** pylandstats.**SpatioTemporalBufferAnalysis**(*landscapes*, *base_mask*, *buffer_dists*, *buffer_rings=False*, *base_mask_crs=None*, *landscape_crs=None*, *landscape_transform=None*, *metrics=None*, *classes=None*, *dates=None*, *metrics_kws={}*)

**__init__**(*landscapes*, *base_mask*, *buffer_dists*, *buffer_rings=False*, *base_mask_crs=None*, *landscape_crs=None*, *landscape_transform=None*, *metrics=None*, *classes=None*, *dates=None*, *metrics_kws={}*)

**Parameters**

- **landscapes** (`list-like`) – A list-like of *Landscape* objects or of strings/file objects/ pathlib.Path objects so that each is passed as the *landscape* argument of *Landscape.__init__*
- **base_mask** (`shapely geometry or geopandas GeoSeries`) – Geometry that will serve as a base mask to buffer around
- **buffer_rings** (`bool, default False`) – If *False*, each buffer zone will consist of the whole region that lies within the respective buffer distance around the base mask. If *True*, buffer zones will take the form of rings around the base mask.
- **base_mask_crs** (`dict, optional`) – The coordinate reference system of the base mask. Required if the base mask is a shapely geometry or a geopandas GeoSeries without the *crs* attribute set
- **landscape_crs** (`dict, optional`) – The coordinate reference system of the landscapes. Required if the passed-in landscapes are *Landscape* objects, ignored if they are paths to GeoTiff rasters that already contain such information.
- **landscape_transform** (`affine.Affine`) – Transformation from pixel coordinates to coordinate reference system. Required if the passed-in landscapes are *Landscape* objects, ignored if they are paths to GeoTiff rasters that already contain such information.
- **metrics** (`list-like, optional`) – A list-like of strings with the names of the metrics that should be computed in the context of this analysis case
- **classes** (`list-like, optional`) – A list-like of ints or strings with the class values that should be considered in the context of this analysis case
- **dates** (`list-like, optional`) – A list-like of ints or strings that label the date of each snapshot of *landscapes* (for DataFrame indices and plot labels)

- **metrics_kws** (*dict, optional*) – Dictionary mapping the keyword arguments (values) that should be passed to each metric method (key), e.g., to exclude the boundary from the computation of *total_edge*, metric_kws should map the string 'total_edge' (method name) to {'count_boundary': False}. The default empty dictionary will compute each metric according to FRAGSTATS defaults.

**property class_metrics_df**
Property that computes the data frame of class-level metrics, which is multi-indexed by the buffer distance, class and date. Once computed, the data frame is cached so further calls to the property just access an attribute and therefore run in constant time.

**property landscape_metrics_df**
Property that computes the data frame of landcape-level metrics, which is multi-indexed by the buffer distance and date. Once computed, the data frame is cached so further calls to the property just access an attribute and therefore run in constant time.

**plot_landscapes**(*cmap=None, legend=True, subplots_kws={}, show_kws={}, subplots_adjust_kws={}*)
Plots each landscape snapshot in a dedicated matplotlib axis by means of the *Landscape.plot_landscape* method of each instance

**Parameters**

- **cmap** (str or *~matplotlib.colors.Colormap*, optional) – A Colormap instance
- **legend** (*bool, optional*) – If `True`, display the legend of the land use/cover color codes
- **subplots_kws** (*dict, optional*) – Keyword arguments to be passed to *plt.subplots*
- **show_kws** (*dict, optional*) – Keyword arguments to be passed to *rasterio.plot.show*
- **subplots_adjust_kws** (*dict, optional*) – Keyword arguments to be passed to *plt.subplots_adjust*

**Returns** **fig** – The figure with its corresponding plots drawn into its axes

**Return type** *matplotlib.figure.Figure*

**plot_metric**(*metric, class_val=None, ax=None, metric_legend=True, metric_label=None, buffer_dist_legend=True, fmt='–o', plot_kws={}, subplots_kws={}*)

**Parameters**

- **metric** (*str*) – A string indicating the name of the metric to plot
- **class_val** (*int, optional*) – If provided, the metric will be plotted at the level of the corresponding class, otherwise it will be plotted at the landscape level
- **ax** (*axis object, optional*) – Plot in given axis; if None creates a new figure
- **metric_legend** (*bool, default True*) – Whether the metric label should be displayed within the plot (as label of the y-axis)
- **metric_label** (*str, optional*) – Label of the y-axis to be displayed if *metric_legend* is *True*. If the provided value is *None*, the label will be taken from the *settings* module
- **buffer_dist_legend** (*bool, default True*) – Whether a legend linking each plotted line to a buffer distance should be displayed within the plot
- **fmt** (*str, default '--o'*) – A format string for *plt.plot*

- **plot_kws** (*dict*) – Keyword arguments to be passed to *plt.plot*
- **subplots_kws** (*dict*) – Keyword arguments to be passed to *plt.subplots*, only if no axis is given (through the *ax* argument)

**Returns** **ax** – Returns the Axes object with the plot drawn onto it

**Return type** axis object

CHAPTER

FIVE

### CHANGE LOG

## 5.1 1.0.2 (25/07/2019)

- fix landscape array dtype in `SpatioTemporalBufferAnalysis`
- included `LICENSE` in `MANIFEST.in`

## 5.2 1.0.1 (24/07/2019)

- deleted Python 2 classifiers in `setup.py`
- fix ValueError message for `landscape_crs` and `landscape_transform` in `BufferAnalysis`
- fix landscape array dtype in `GradientAnalysis` and `BufferAnalysis`

## 5.3 1.0.0 (18/07/2019)

- dropped Python 2 support
- added `SpatioTemporalBufferAnalysis.plot_landscapes` method
- added `buffer_dist_legend` argument and docs in `SpatioTemporalBufferAnalysis.plot_metric`
- fix proper metric data frame properties in `SpatioTemporalBufferAnalysis`
- pass `transform` argument when initializating `MultiLandscape` instances (i.e., `SpatioTemporalAnalysis`, `BufferAnalysis`, `GradientAnalysis` and `SpatioTemporalBufferAnalysis`)
- `plot_landscape` and `plot_landscapes` with rasterio.plot.show
- changed `feature_{name,values}` for `attribute_{name,values}` in `MultiLandscape` (abstract) class
- dropped `plot_metrics` method

## 5.4 0.6.1 (02/07/2019)

- cell width and length comparisons with `numpy.isclose` to deal with imprecisions that come from float pixel resolutions (e.g., in GeoTIFF files)

- critical fix: moved pythran signatures from `compute.pythran` to `compute.py` so that the build is properly done when pip-installing

## 5.5 0.6.0 (01/07/2019)

- flat array approach to the computation of the adjacency matrix with pythran to improve performance (plus fixes a bug on the computation of `total_edge`)
- initialization of `scipy.spatial.cKDTree` with keyword arguments `balanced_tree` and `compact_nodes` set to `False`

## 5.6 0.5.0 (28/05/2019)

- methods plotting multiple axes return only the figure instead of a tuple with the figure and the axes
- settings module that allow configuring metrics' labels in plots, defaulting to FRAGSTATS abbreviations
- chaged CRS in tests to work with pyproj >= 2.0.0
- corrected `figlength` for `figwidth`
- warn when computing metrics that require an unmet minimum number of patches or classes (and their computation returns nan)
- all tests with `unittest.TestCase` assert methods

## 5.7 0.4.1 (03/04/2019)

- added docstrings for `MultiLandscape`, `GradientAnalysis` and `BufferAnalysis`
- raise `ValueError` when using buffer rings around a polygon geometry

## 5.8 0.4.0 (03/04/2019)

- implemented `SpatioTemporalBufferAnalysis` with a dedicated `plot_metric` method
- added buffer ring-wise analysis through a `buffer_rings` boolean argument in `BufferAnalysis`

## 5.9 0.3.1 (29/03/2019)

- float equality comparations with numpy `isclose` method

## 5.10 0.3.0 (28/03/2019)

- implemented `GradientAnalysis` and `BufferAnalysis`
- added optional `geopandas` dependences
- created abstract `MultiLandscape` class

- `Landscape` initialization from ndarray or geotiff (dropped `read_geotiff` method)
- implemented `contagion`
- convolution-based adjacency dataframe
- fixed bug with `class_cond` in `Landscape.compute_arr_edge`

## 5.11 0.2.0 (18/03/2019)

- implemented `euclidean_nearest_neighbor` and all its corresponding class/landscape distribution statistic metrics
- fixed dtype of `self.classes` in `landscape.Landscape`
- set default argument `nodata=None` (instead of `nodata=0`) for `landscape.read_geotiff`
- changed test input data files
- implemented `__len__` of `spatiotemporal.SpatioTemporalAnalysis`

## 5.12 0.1.1 (22/01/2019)

- corrected if-else statements involving `None` variables

## 5.13 0.1.0 (22/01/2019)

- initial release

CHAPTER

SIX

### CONTRIBUTING

Contributions are always greatly appreciated and credit will always be given.

#### 6.1 Types of contributions

##### 6.1.1 Report bugs

Report bugs at https://github.com/martibosch/pylandstats/issues.

If you are reporting a bug, please include:

- Your operating system name and version.
- Any details about your local setup that might be helpful in troubleshooting.
- Detailed steps to reproduce the bug.

##### 6.1.2 Fix bugs

Look through the GitHub issues for bugs. Anything tagged with "bug" and "help wanted" is open to whoever wants to implement it.

##### 6.1.3 Implement features

Look through the GitHub issues for features. Anything tagged with "enhancement" and "help wanted" is open to whoever wants to implement it.

#### 6.2 Pull request guidelines

Before you submit a pull request, check that it meets these guidelines:

1. The pull request should include tests.
2. If the pull request adds functionality, the docs should be updated. Put your new functionality into a function with a docstring, and add the feature to the list in README.md.
3. The pull request should work for Python 2.7 and 3.6, as well as for PyPy. Check https://travis-ci.org/martibosch/pylandstats/pull_requests and make sure that the tests pass for all supported Python versions.

This documentation is intended as an API reference. See also:

- pylandstats-notebooks repository (tutorial/thorough overview of PyLandStats)
- swiss-urbanization repository (example application of PyLandStats to evaluate the spatiotemporal patterns of urbanization in three Swiss urban agglomerations)

CHAPTER

SEVEN

### FEATURES

- Compute pandas DataFrames of landscape metrics at the patch, class and landscape level
- Analyze the spatiotemporal evolution of landscapes
- Analyze landscape changes accross environmental gradients

CHAPTER

EIGHT

### USING PYLANDSTATS

To install use pip:

```
$ pip install pylandstats
```

If you want to use the `BufferAnalysis` class, you will need geopandas. The easiest way to install it is via conda-forge as in:

```
$ conda install -c conda-forge geopandas
```

and then install PyLandStats with the `geo` extras as in:

```
$ pip install pylandstats[geo]
```

CHAPTER

NINE

### INDICES AND TABLES

- genindex
- modindex

### INDEX

#### Symbols


__init__() (*pylandstats.BufferAnalysis method*), 14
__init__() (*pylandstats.GradientAnalysis method*), 13
__init__() (*pylandstats.SpatioTemporalAnalysis method*), 11
__init__() (*pylandstats.SpatioTemporalBufferAnalysis method*), 17


## A


area() (*pylandstats.Landscape method*), 3


## B


BufferAnalysis (*class in pylandstats*), 14


## C


class_metrics_df() (*pylandstats.BufferAnalysis property*), 15
class_metrics_df() (*pylandstats.GradientAnalysis property*), 13
class_metrics_df() (*pylandstats.SpatioTemporalAnalysis property*), 11
class_metrics_df() (*pylandstats.SpatioTemporalBufferAnalysis property*), 18
compute_class_metrics_df() (*pylandstats.Landscape method*), 9
compute_landscape_metrics_df() (*pylandstats.Landscape method*), 9
compute_patch_metrics_df() (*pylandstats.Landscape method*), 9
contagion() (*pylandstats.Landscape method*), 8


## E


edge_density() (*pylandstats.Landscape method*), 7
euclidean_nearest_neighbor() (*pylandstats.Landscape method*), 5


## F


fractal_dimension() (*pylandstats.Landscape method*), 4


## G


GradientAnalysis (*class in pylandstats*), 13


## L


landscape_metrics_df() (*pylandstats.BufferAnalysis property*), 15
landscape_metrics_df() (*pylandstats.GradientAnalysis property*), 13
landscape_metrics_df() (*pylandstats.SpatioTemporalAnalysis property*), 11
landscape_metrics_df() (*pylandstats.SpatioTemporalBufferAnalysis property*), 18
landscape_shape_index() (*pylandstats.Landscape method*), 8
largest_patch_index() (*pylandstats.Landscape method*), 6


## N


number_of_patches() (*pylandstats.Landscape method*), 6


## P


patch_density() (*pylandstats.Landscape method*), 6
perimeter() (*pylandstats.Landscape method*), 4
perimeter_area_ratio() (*pylandstats.Landscape method*), 4
plot_landscape() (*pylandstats.Landscape method*), 10
plot_landscapes() (*pylandstats.BufferAnalysis method*), 15
plot_landscapes() (*pylandstats.GradientAnalysis method*), 13
plot_landscapes() (*pylandstats.SpatioTemporalAnalysis method*), 11

plot_landscapes() (*pylandstats.SpatioTemporalBufferAnalysis method*), 18
plot_metric() (*pylandstats.BufferAnalysis method*), 15
plot_metric() (*pylandstats.GradientAnalysis method*), 14
plot_metric() (*pylandstats.SpatioTemporalAnalysis method*), 12
plot_metric() (*pylandstats.SpatioTemporalBufferAnalysis method*), 18
proportion_of_landscape() (*pylandstats.Landscape method*), 5

## S

shannon_diversity_index() (*pylandstats.Landscape method*), 8
shape_index() (*pylandstats.Landscape method*), 4
SpatioTemporalAnalysis (*class in pylandstats*), 11
SpatioTemporalBufferAnalysis (*class in pylandstats*), 17

## T

total_area() (*pylandstats.Landscape method*), 5
total_edge() (*pylandstats.Landscape method*), 7
